# Elongation of the nascent avian foregut requires coordination of intrinsic and extrinsic cell behaviors

**DOI:** 10.1101/2024.10.31.621372

**Authors:** Olivia Powell, Emily Garcia, Vanshika Sriram, Yi Qu, Nandan L. Nerurkar

**Affiliations:** Department of Biomedical Engineering, Columbia University, New York NY 10027

## Abstract

The foregut tube gives rise to the lungs and upper gastrointestinal tract, enabling vital functions of respiration and digestion. How the foregut tube forms during embryonic development has historically received considerable attention, but over the past few decades this question has primarily been addressed indirectly through studies on morphogenesis of the primitive heart tube, a closely related process. As a result, many aspects of foregut development remain unresolved. Here, we exploit the accessibility of the chick embryo to study the initial formation of the foregut tube, combining embryology with fate mapping, live imaging, and biomechanical analyses. The present study reveals that the foregut forms and elongates over a narrower time window than previously thought, and displays marked dorso-ventral and left-right asymmetries early in its development. Through tissue-specific ablation of endoderm along the anterior intestinal portal, we confirm its central biomechanical role in driving foregut morphogenesis, despite not directly contributing cells to the elongating tube. We further confirm the important role of this cell population in formation of the heart tube, with evidence that this role extends to later stages of cardiac looping as well. Together, these data reveal the need for an intricate balance between intrinsic cell behaviors and extrinsic forces for normal foregut elongation, and set the stage for future studies aimed at understanding the underlying molecular cues that coordinate this balance.

**GRAPHICAL ABSTRACT:** 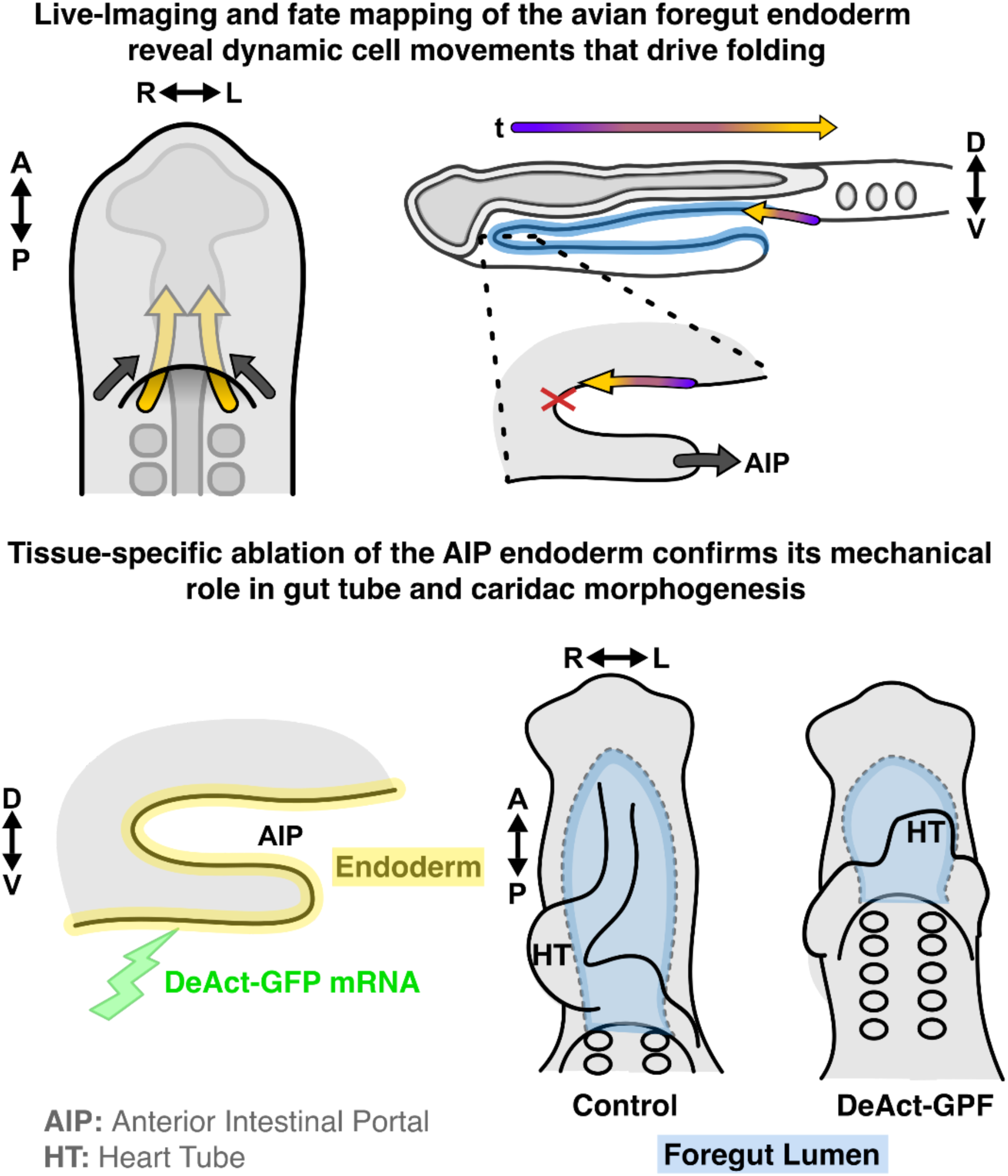

## 1. Introduction

The respiratory and digestive organs arise during embryonic development from a simple epithelial cylinder, the gut tube. In turn, the gut tube forms from an initially flat endodermal epithelium that undergoes dramatic physical transformations as it internalizes from the ventral surface of the embryo, a process initiating shortly after the onset of gastrulation. Despite the conserved function of gut-derived organs, the macroscopic process by which the tube forms is divergent between amniotes and anamniotes. Among amniotes, the gut tube forms as three distinct segments along the antero-posterior axis: the foregut, midgut, and hindgut. Remarkably, while they form a single continuous tube, the timing, location, and mechanism of tube formation differs for each segment (Durel and Nerurkar 2020). While historically, the question of how the gut tube forms has received considerable attention (Wolff 1769; Bellairs 1953; Stalsberg and DeHaan 1968; Rosenquist 1971; Seidl and Steding 1978), work over the past two to three decades has largely considered amniote foregut morphogenesis primarily as it relates to heart development. The AIP endoderm has been suggested to act as a key organizing center for differentiation of precardiac mesoderm (Schultheiss, Xydas, and Lassar 1995; Anderson et al. 2016), and to guide formation of the primitive heart tube (Kirby et al. 2003; Varner and Taber 2012; Aleksandrova et al. 2012; Hosseini, Garcia, and Taber 2017; Kidokoro et al. 2018). As a result, many outstanding questions remain regarding the initiation and elongation of the foregut tube.

In the chick embryo, foregut formation begins with a crescent shaped invagination of the endoderm, known as the anterior intestinal portal (AIP), which is thought to fold passively due to forces generated in the neighboring neural plate during head-fold formation at Hamburger Hamilton (HH) stages 6 to 7 (Hamburger and Hamilton 1951; Varner, Voronov, and Taber 2010; Bellairs and Osmond 2014). With its formation, the AIP creates a shallow pocket of endoderm on the ventral surface, which progressively deepens into a blind-ended tube as the AIP descends posteriorly during development (Figure 1A). Throughout this process, the AIP lip forms the physical boundary between the internalized foregut endoderm and extraembryonic ventral epithelium that remains exposed to the yolk. As the AIP descends, it is also thought to draw together the bilateral heart fields, which fuse at the midline to form the primitive heart tube. Movement of precardiac mesoderm closely mirrors tissue movements in the endoderm along the AIP lip (Cui et al. 2009; Aleksandrova et al. 2012; Kidokoro et al. 2018). Biomechanical investigations suggest that cytoskeletal contraction within this region may provide the driving force, actively pulling the bilateral heart fields along until they meet at the embryonic midline (Varner and Taber 2012; Hack et al. 2023). These same contractions along the lip of the AIP have been proposed to drive descensus of the AIP and elongation of the foregut (Hosseini, Garcia, and Taber 2017). However, evidence for this is primarily based on broad pharmacologic perturbations. As a result, in the absence of tissue-specific perturbations that directly target the endoderm, the biomechanical importance of AIP endoderm for heart tube fusion and foregut elongation has not been directly tested. Moreover, because the AIP is externally accessible, much of the efforts on foregut tube morphogenesis have focused on descensus of the AIP as a direct correlate of foregut tube formation (Bellairs 1953; Stalsberg and DeHaan 1968; Varner and Taber 2012; Hosseini, Garcia, and Taber 2017), without directly examining links between the two. As a result, much remains unresolved regarding the tissue-scale mechanisms of foregut elongation. In particular, it remains unclear whether elongation of the gut requires intrinsic cell behaviors or can be largely reduced to movement of the AIP, as it has been primarily discussed in the past. The present study combines embryological approaches with live imaging and biomechanical analyses in the chick embryo to provide new insights into the cellular basis of foregut morphogenesis. Based on embryological measurements, we concluded that AIP descensus is in fact not a direct indication of foregut elongation, and through a series of additional experiments reframe the question of foregut morphogenesis as a balancing act between intrinsic and extrinsic cell behaviors that coordinate this complex, three-dimensional process.

**Figure 1:**
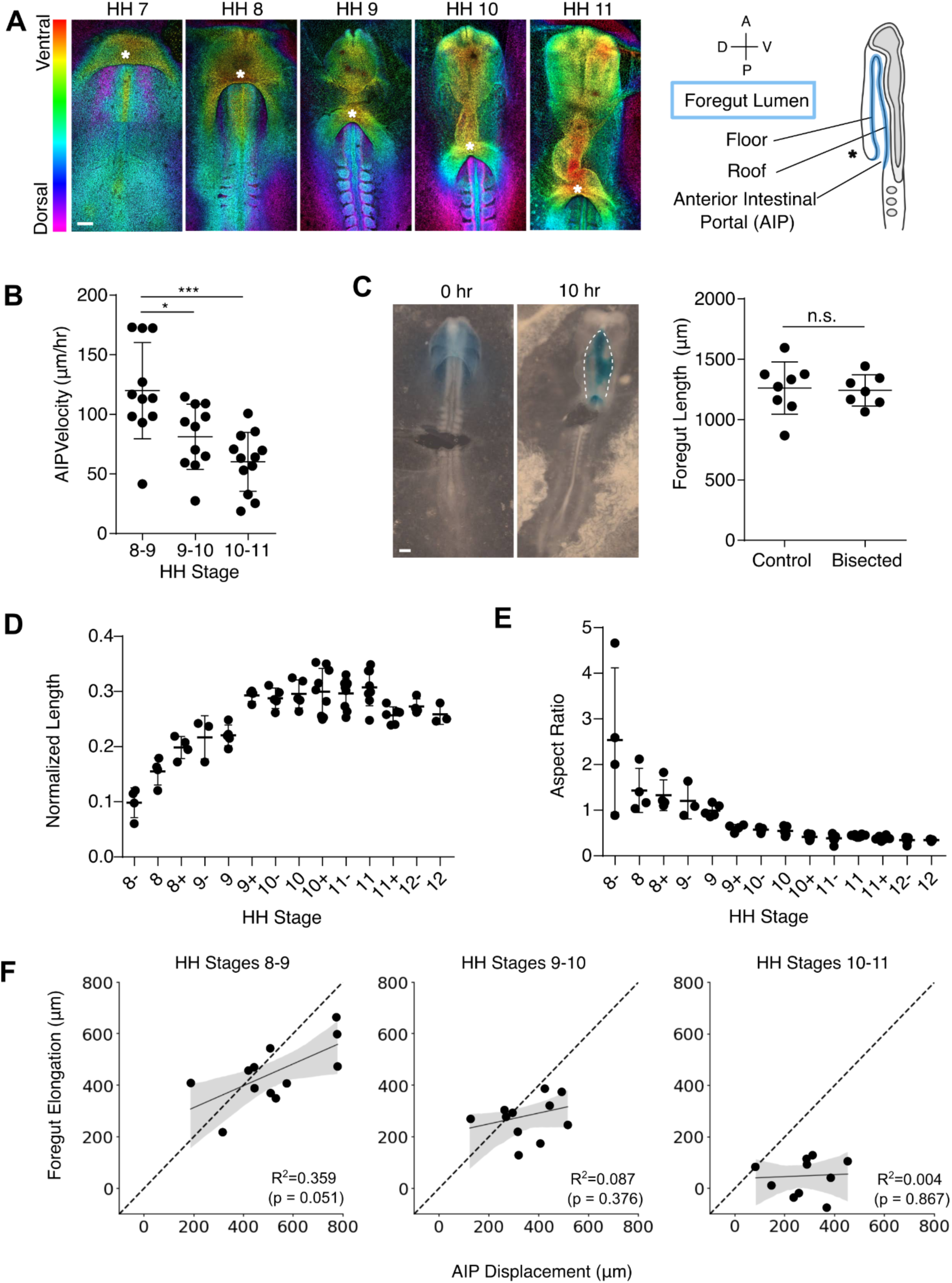
AIP displacement is poorly correlated with foregut elongation. (**A,** left) Representative depth-coded maximum intensity projections of DAPI-stained chick embryos from the initiation of foregut formation with the emergence of the head fold at HH stage 7 to onset of c-looping of the heart tube at HH stage 11. *indicates the position of the medial Anterior Intestinal Portal (AIP); scale = 200 μm. (**A**, right) Schematic sagittal section of the anterior chick embryo during foregut elongation. The forming foregut lumen is highlighted in blue, with the ventral foregut floor and dorsal foregut roof indicated. (**B**) AIP velocity between HH stages 8-9, 9-10, and 10-11 (n = 10 per group); *p<0.05, ***p<0.001 as determined by one-way ANOVA with Tukey’s multiple comparisons test. (**C**, left) HH stage 8 embryos following bisection (0 hours) to mechanically separate AIP from regressing node region and following 10 hours incubation ; scale = 200 µm. (**C**, right) Foregut length compared between bisected (n = 7) and intact (n = 8) control embryos; n.s. = not significant (p = 0.211 by Student’s t-test). (**D, E**) Foregut tube length normalized to antero-posterior embryo length (**D**) and foregut aspect ratio (**E**) across developmental stages HH stage 8- to 12 (n = 4-7 per stage). (**F**) Comparison of foregut elongation versus AIP displacement of individual embryos (n = 10 per stage) between HH stages 8-9 (left), 9-10 (middle), and 10-11 (right). Linear regression is indicated by a solid line, with shaded regions representing the 95% confidence interval calculated for the regression estimate, and dotted lines indicate y = x. Pearson correlation coefficient was used to measure the linear relationship between the two measures.

## 2. Results

### 2.1 Dynamics of AIP displacement do not depend on axis elongation

Guided by prior studies on foregut morphogenesis, which have primarily focused on the AIP, we began our efforts to understand the mechanisms of foregut tube morphogenesis by investigating the posterior-ward movement of the AIP (Figure 1A). Based on correlation between the rates and AIP displacement and regression of Hensen’s node, it has previously been proposed that movement of the AIP is a passive result of forces generated by axis elongation (Seidl and Steding 1978). Although more recent studies have focused on the role of contractile forces within the AIP region itself (Varner and Taber 2012;Shi, Varner, and Taber 2015; Kidokoro et al. 2018; Hack et al. 2023), a role for axis elongation in pulling the AIP posteriorly has not been directly tested. To do so, we first quantified AIP movement over time. Prior studies have quantified AIP displacement by measuring the change in distance between the AIP and the anterior neuropore (Stalsberg and DeHaan 1968; Hosseini, Garcia, and Taber 2017), likely overestimating AIP movement due to anterior displacement of the neuropore itself (Seidl and Steding 1978). Using somite 2 (which remains relatively stationary, Supplemental Figure S1A, B), as a reference point, we found that AIP descensus slows over time between HH stages 8 and 11 (Figure 1B), even as axis elongation proceeds at a constant rate (Bénazéraf et al. 2010; Xiong et al. 2020). It is therefore unlikely that AIP movement is simply due to pulling forces from the regressing Henen’s node. To test this further, we bisected embryos along the antero-posterior axis at HH stage 8 to mechanically decouple posterior elongation of the embryo from morphogenesis of the foregut tube. Anterior and posterior segments of bisected embryos developed normally, and measurement of foregut length by injection of Fast Green dye into the lumen revealed no significant difference when compared to intact control embryos (Figure 1C, Supplementary Figure S1C). These results indicate that axis elongation is not required for posterior movement of the AIP, and are consistent with the view that local forces may be the driver of AIP movement during foregut formation (Shi, Varner, and Taber 2015; Hosseini, Garcia, and Taber 2017).

### 2.2 AIP displacement is a poor indicator of foregut tube elongation

Prior studies of foregut morphogenesis have largely equated elongation of the nascent foregut tube with posterior displacement of the AIP. However, to what extent the movement of this external structure reflects true elongation of the foregut as an epithelial tube internally has not been investigated. To distinguish between AIP movement and elongation of the foregut across developmental stages, we measured tube length and aspect ratio directly via injection of Fast Green through the AIP (Supplemental Figure S1C). This revealed that foregut elongation initially proceeds at a constant rate between HH stages 8- and 10- (Figure 1D). Surprisingly, however, gut length plateaus from HH stage 10 onward (Figure 1D), despite the continued descensus of the AIP (Figure 1A, B). Elongation of the foregut tube is accompanied by medio-lateral narrowing, which produces a corresponding decrease in the aspect ratio of the tube (Figure 1E). To directly test the correlation between AIP descensus and foregut length, we measured both in the same embryos over time. This revealed that the two measures are only weakly correlated during the initial stages of foregut elongation, and as development proceeds, even this weak correlation is lost (Figure 1F). These findings suggest that it is not possible to draw conclusions on the events driving foregut tube elongation by observing movement of the AIP alone.

### 2.3 Volumetric imaging of the nascent foregut tube reveals intrinsic remodeling and early emergence of left-right asymmetry

Based on the surprising observation that AIP descensus continues after foregut elongation plateaus (Figure 1), we next asked whether AIP movement may incorporate tissue into the forming foregut through increasing surface area while maintaining tube length. To do so, 3-D morphology of the foregut tube was reconstructed from confocal imaging of optically cleared samples stained with DAPI to visualize nuclei. Throughout its development, the nascent foregut tube remains remarkably flat along the dorso-ventral axis, with a slight dorsal bend laterally, and a lumen that much less resembles a tube than an envelope: apical surfaces of foregut floor and roof (ventral and dorsal, respectively, as indicated in Figure 1A) are separated by a lumen of approximately 20-30 μm in width (Figure 2A). As development proceeds, the foregut cross-section becomes increasingly non uniform along the antero-posterior axis, with regional constrictions appearing to segment the foregut by HH stage 12. Surprisingly, we observed an emergent dilation of the right lateral boundary of the foregut tube at HH stages 11 to 12 (Figure 2A), preceding - to our knowledge - any prior reports of left-right asymmetry in the gut (Davis et al. 2008; Grzymkowski, Wyatt, and Nascone-Yoder 2020). Together, these results suggest that the groundwork for later organogenesis of foregut-derived organs is likely underway early, with complex reorganization of the foregut occurring as it forms. Quantification of luminal volume and surface area of the foregut revealed a similar dynamic to elongation, with both increasing between HH stages 8 and 10, and no significant changes from HH stage 10 and 12 (Figure 2B, C). Despite the remarkable changes in foregut size and shape that occur between HH stages 8 and 12, the surface area-to-volume ratio of the foregut tube remained surprisingly constant over this period (Figure 2D). Taken together, these data suggest that after HH stage 10, further descensus of the AIP may reflect posterior translation of the foregut tube itself, rather than elongation or expansion of the structure. As a result, elongation of the nascent foregut occurs during a narrower window than previously thought, between HH stages 8 and 10-. Moving forward, we focus on these stages to ask how intrinsic and extrinsic processes contribute to elongation of the foregut.

**Figure 2:**
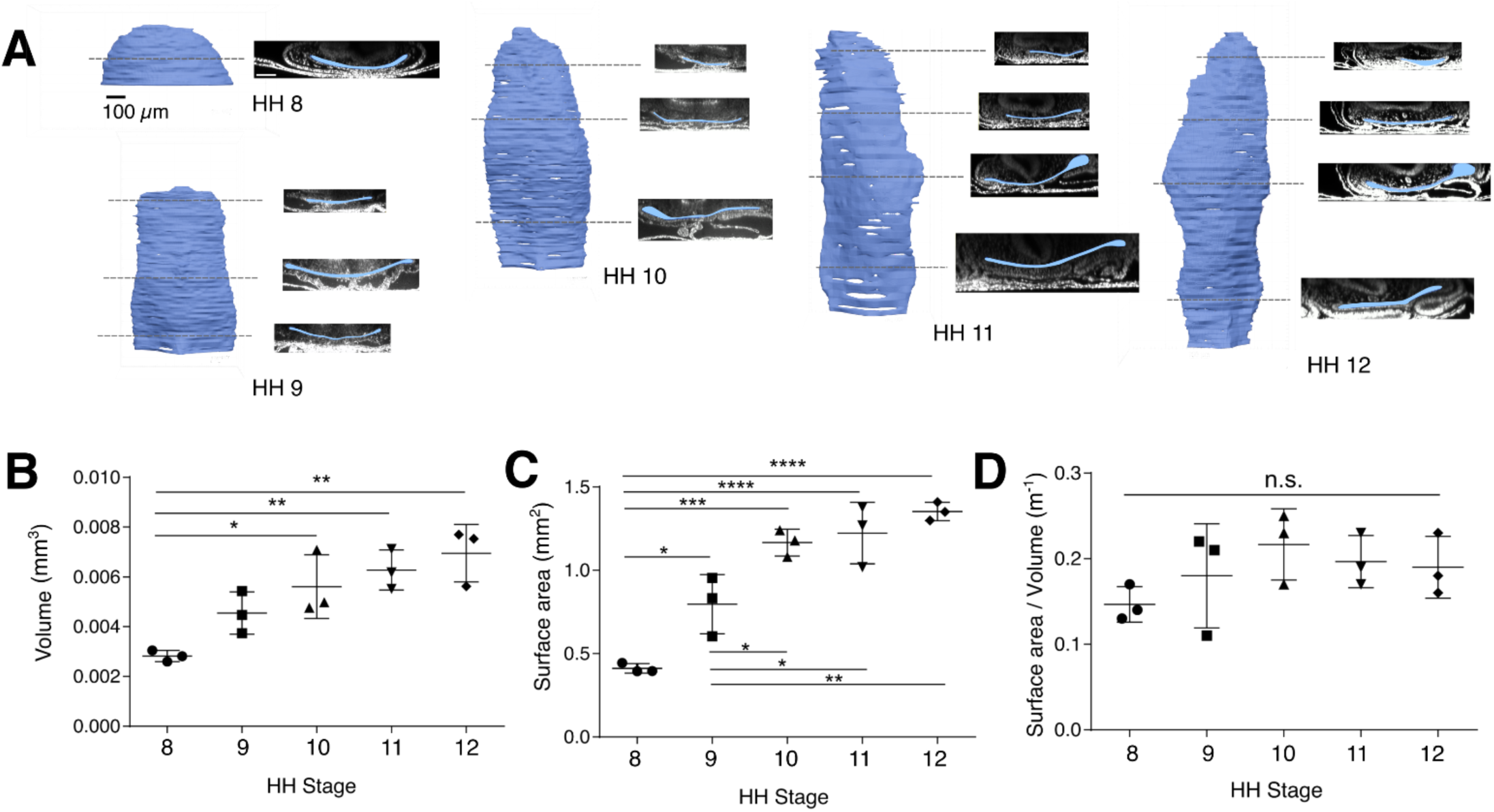
3D volumetric imaging of the nascent foregut lumen. (**A**) At each HH stage between 8 and 12, 3-D reconstructions of the foregut lumen volume are shown at left, with transverse optical sections along the antero-posterior axis shown at right (lumen pseudocolored blue); scale bar = 100 µm. (**B**-**D**) Time course quantification of lumen volume (**B**), surface area (**C**) , and surface area-to-volume ratio (**D**) between HH stages 8 and 12; *p<0.05, **p<0.01, ***p<0.001 as determined by one-way ANOVA with Tukey’s multiple comparisons test (n = 3 per stage).

### 2.4 Roll-over of cells contributes minimally to elongation of the foregut tube

To further investigate the relationship between AIP movement and foregut elongation, we next considered movement of endoderm cells relative to the AIP as it moves posteriorly. There are conflicting reports in the literature as to whether “in-rolling” of cells occurs across the AIP lip into the forming foregut (Bellairs 1953; Stalsberg and DeHaan 1968). These studies, carried out decades ago, relied on ferric oxide or carbon particles to label tissues and examine their movements. We therefore sought to first resolve this question using more precise current approaches, marking cells by microinjection of the fluorescent, lipophilic dye DiI. Displacement of DiI-labeled cells along the AIP was examined in 4 hour increments between HH stage 7 and HH stage 10. During these stages, when foregut length increases by 3-fold (Figure 1D), labeled cells largely remained on the ventral surface of the AIP as it moved, with little to no in-rolling over the AIP lip (Figure 3A). This raises the question of how the midline foregut floor accommodates elongation strains of over 200%, if material is not added from cells crossing the AIP boundary to incorporate into the foregut floor. In previous work, we determined that the hindgut elongates by inversion of cells from the hindgut roof to the floor by passing over the blind end of the tube (Nerurkar et al. 2019). We therefore asked whether a similar process may occur in the foregut to enable elongation of the floor by rolling over of cells at the blind anterior end of the tube. To test this, we injected lipophilic vital dyes, DiI and DiO, to label two antero-posterior positions along the midline of the presumptive roof endoderm (Figure 3B), and asked whether they invert across the anterior end of the foregut as it forms. Populations marked by injection of DiI and DiO at HH stage 8- did not invert by stage 10, and remained restricted to the roof (n = 4/4). This suggests that foregut floor elongation cannot be explained by rolling over of roof cells at the anterior end of the elongating foregut, pointing instead to cell behaviors intrinsic to the foregut floor as the driver of tissue elongation. Notably, however, while the two labeled populations remained in the roof, they did converge anteriorly toward the blind end of the tube. Therefore, while there may be considerable cell rearrangements within roof and floor compartments of the forming foregut, there is little to no mixing between these tissues along the midline.

**Figure 3:**
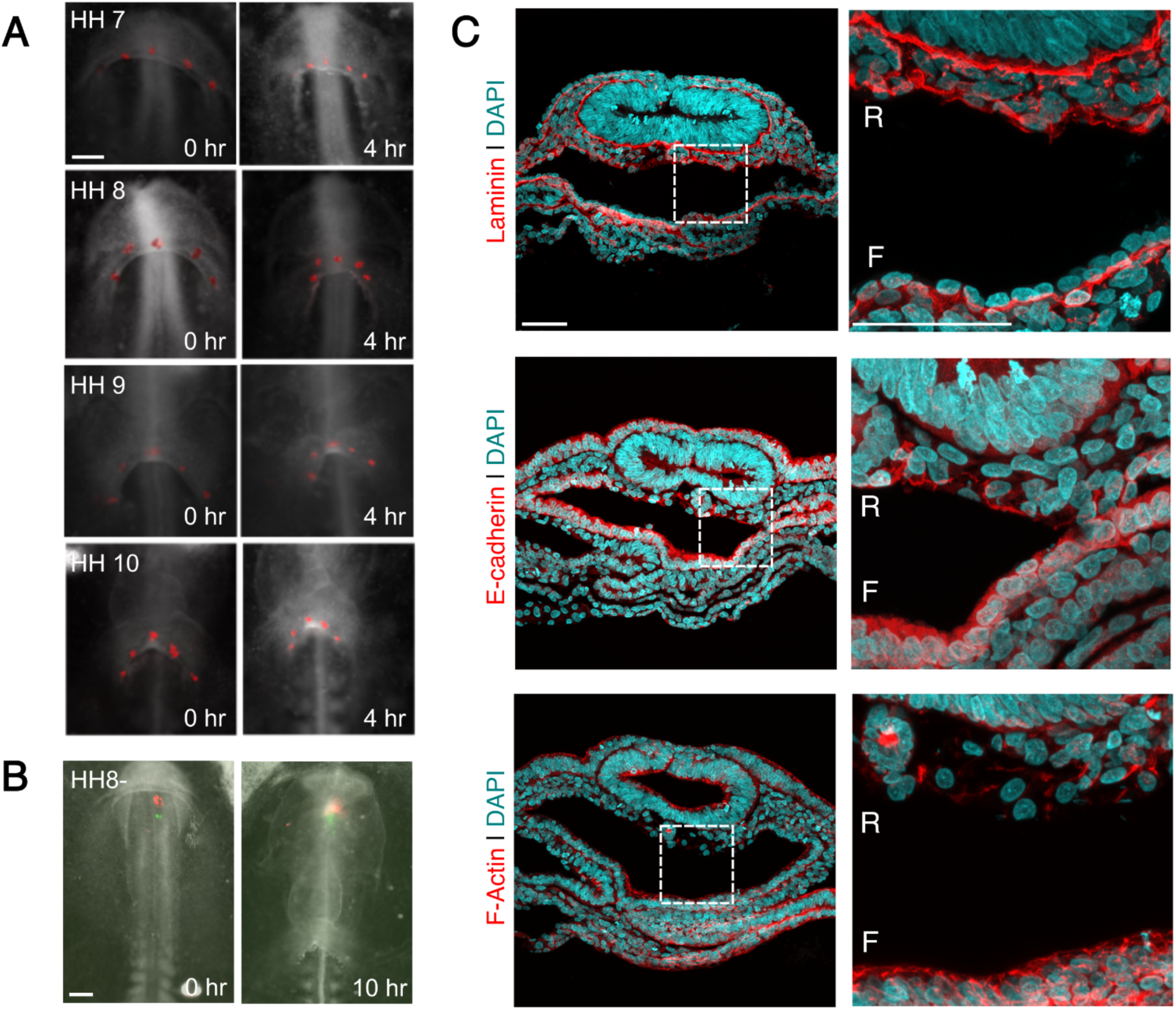
Fate maps reveal limited involution of the midline endoderm across the AIP. (**A**) Di labels injected onto AIP endoderm between HH stages 7-10 (left, 0 hours) and following 4 hours incubation (right); 4 hours spans the approximate time between HH stages in this range (HH7, n = 4; HH8, n = 7; HH9, n = 5; HH10, n = 3); scale = 200 µm. (**B**) DiI (red) and DiO (green) labels injected into midline presumptive roof endoderm at HH stage 8- move anteriorly toward the prechordal plate, but remain on the roof at HH stage 10+ (n = 4/4); scale = 200 µm. (**C**) Transverse sections through the foregut at HH stage 10 immunostained (red) for laminin (top) and E-cadherin (middle), and stained with phalloidin to visualize F-actin (bottom); counterstained with DAPI to visualize nuclei (cyan); R = foregut roof; F = foregut floor; scale = 50 μm (n = 3/3).

### 2.5 The forming foregut displays early dorso-ventral asymmetry of endoderm cells

Above, we observed that roof and floor cells of the foregut midline do not mix during elongation of the foregut tube (Figure 3B), despite considerable reorganization of cells within the roof. To examine this further, we histologically characterized cells of the forming foregut roof and floor to examine potential phenotypic disparities. Visualization of epithelial markers revealed striking differences between the two populations. The foregut floor displayed several hallmarks of a columnar epithelium, including a well-formed laminin-rich basement membrane located at the basal surface of the endoderm, and apical and basolateral enrichment of E-Cadherin and F-actin (Figure 3C). By contrast, the foregut roof was more loosely organized with lower cell density, flatter cells lacking clear apico-basal polarity, and a disorganized basement membrane that ensconced individual cells rather than forming a single continuous basal layer (Figure 3C). These results suggest that at the same antero-posterior level, roof and floor endoderm cells represent distinct cell populations based on their cellular organization. Although differences in epithelial thickness between the roof and floor have been previously recognized (Adelmann 1922; Bellairs 1953), these results suggest that foregut roof endoderm adopts an almost mesenchymal phenotype compared to the more conventional columnar epithelium of the foregut floor. Further, these findings indicate that dorso-ventral symmetry of the gut tube is broken as the foregut forms (Lyons, Hogan, and Robertson 1995; Zorn and Wells 2009; Ioannides et al. 2010; Schäfer et al. 2021).

### 2.6 Tissue-specific ablation of AIP endoderm disrupts foregut elongation and cardiac morphogenesis

Acto-myosin activity of endoderm cells along the AIP lip is thought to drive descensus of the AIP and elongation of the foregut tube (Aleksandrova et al. 2012; Varner and Taber 2012; Shi, Varner, and Taber 2015; Hosseini, Garcia, and Taber 2017; Kidokoro et al. 2018). However, the primary support for this comes from descriptive studies of cell shape and myosin organization, quantification of tissue deformations that are consistent with contractility (Supplemental Figure S2), or from broad pharmacologic perturbations that disrupt acto-myosin activity throughout the entire embryo. As a result, while many observations are consistent with the hypothesis that the AIP endoderm plays a central mechanical role in foregut elongation, it has not been directly tested. To do so, we relied on tissue-specific transfection of the endodermal epithelium (Nerurkar et al. 2019) to test whether endoderm-specific perturbations of contractility in the AIP influence foregut elongation. Specifically, we made use of DeAct-SpvB, a genetically encoded ADP ribosyltransferase that severely disrupts F-actin (Harterink et al. 2017; Oikonomou et al. 2024) fused to GFP (referred to below as DeAct-GFP), to exclusively remove acto-myosin activity in the endoderm. To greatly increase efficiency of misexpression along the AIP, RNA electroporation was used in place of plasmid DNA (Tran, Dave, and Lansford 2019). This produced widespread disruption of F-actin throughout the endoderm (Figure 4A). In fact, these effects were so pronounced that in many cases, electroporated endoderm cells delaminated from the embryo, leaving behind an intact basement membrane supporting a discontinuous endodermal epithelium (Supplemental Figure S3). As a result, these experiments effectively investigate the effects of endoderm ablation, rather than solely disruption of endoderm contractility. DeAct misexpression at HH stage 8 caused a significant reduction in foregut elongation and a broadening of the AIP (Figure 4B, C), suggesting that the endodermal layer at the AIP is required for elongation of the foregut, despite these cells not contributing directly to the foregut tube. This is consistent with AIP endoderm generating local tension to drive descensus of the AIP, facilitating elongation of the foregut through intrinsic mechanisms. It is notable that upon ablation of endoderm AIP, the foregut did continue to elongate by nearly 50%, suggesting that there may be some contributions to AIP elongation that are not mechanically dependent on the AIP endoderm.

**Figure 4:**
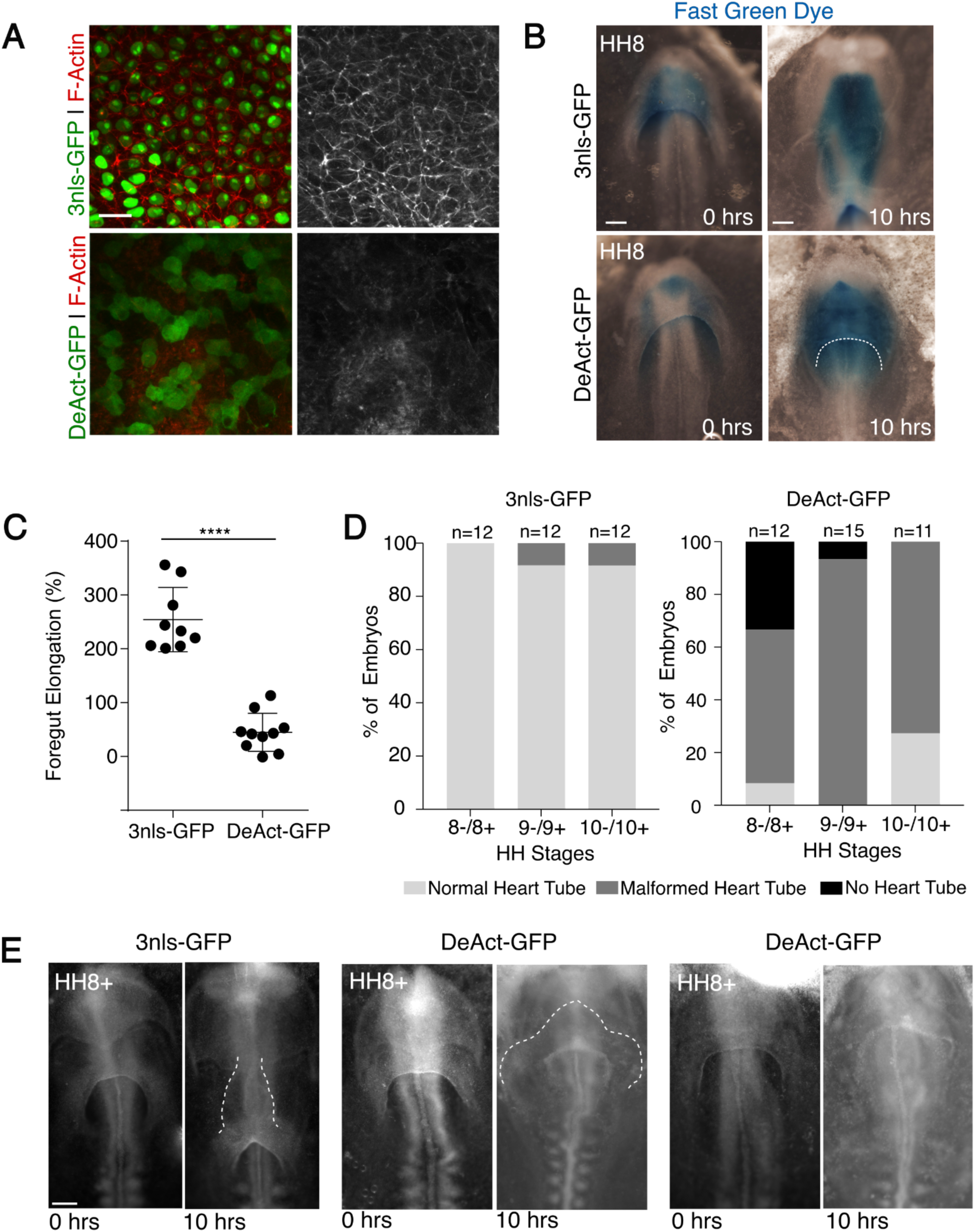
Role of endoderm contractility in AIP movement and FG formation. (**A**) mRNA electroporation 3nls-GFP and DeAct-GFP into AIP endoderm of HH stage 10 embryos followed by phalloidin staining to visualize effects on F-actin integrity. Phalloidin signal isolated in gray scale at right; scale = 20 μm. (**B**) Fast Green injection into the foregut lumen to visualize foregut length (quantified in **C**) immediately following electroporation with 3nl-GFP or DeAct-GFP mRNA at HH stage 8, and again after 10 hours incubation. Dashed white line indicates AIP lip; scale = 200 μm. (**C**) Percent elongation of foregut following mRNA electroporation of AIP endoderm with 3nls-GFP (n=9) or DeAct-GFP (n=10); ****p < 0.0001 by Student’s t-test. (**D**) Percentage of embryos displaying heart tube defects with 10 hours incubation following mRNA electroporation with 3nls-GFP (left) or DeAct-GFP mRNA (right). (**E**) Representative images of embryos electroporated at HH stage 8+ with 3nls-GFP, yielding normal control embryos (left), compared to DeAct-GFP electroporated embryos that displayed either heart tube malformations (middle) no heart tube at all (right). White dashed lines indicate heart tube boundary; scale = 200 μm.

Prior studies suggest that endoderm contraction along the AIP lip is responsible for pulling the bilateral heart fields toward the midline and driving their fusion to form the heart tube (Varner and Taber 2012; Aleksandrova et al. 2012). We next tested this by performing endoderm ablations using the DeAct-GFP electroporations targeting the AIP endoderm at a range of stages between HH stage 8-, when AIP descensus begins, and HH10+, when foregut length has plateaued and the linear heart tube is fully formed. Embryos were incubated *ex ovo* for 10 hours to examine the effects on heart morphogenesis. Endoderm ablation dramatically disrupted normal cardiac morphogenesis across all stages, producing phenotypes with severity ranging from complete agenesis of the heart tube to formation of an abnormally looped heat tube, with a small subset of embryos producing normally looped hearts as well (Figure 4D, E and Supplemental Figure S4). Although endoderm ablation at later stages had a somewhat diminished effect, even at HH stage 10 when the process of heart tube fusion is nearly complete, DeAct-GFP electroporation disrupted normal dextral looping of the heart tube into a C-shaped bend (c-looping) in over 70% of embryos (Figure 4D, E). These data directly confirm the importance of endoderm for migration of the bilateral heart fields and morphogenesis of the primitive heart tube. In addition, looping abnormalities after misexpression of DeAct-GFP at HH10+ suggest that the endoderm plays a previously unappreciated role in cardiac c-looping, a fundamental step in transforming the primitive heart tube into a four chambered pump.

### 2.7 Live in vivo imaging of cell movements reveals rapid polarization along the AIP and dynamic tissue deformations during AIP descensus

Because gross measures of AIP displacement proved to be a poor indicator of foregut elongation (Figure 1F), we asked how cell movements in the endoderm lining the AIP relate to macroscopic movement of the AIP itself. While tissue displacements along the AIP have been visualized by observing movement of adherent particles (Bellairs 1953) or sparsely applied vital dyes (Varner and Taber 2012; Shi, Varner, and Taber 2015; Hack et al. 2023), it has been challenging to link these tissue movements to underlying cell behaviors because live cell imaging of the forming foregut endoderm has been limited (Ivanovitch, Temiño, and Torres 2017). We therefore set out to visualize endoderm cell dynamics during descensus of the AIP and formation of the foregut. Embryos were subject to mosaic electroporation with a ubiquitous GFP reporter to enable live imaging of cell morphology and movements throughout foregut tube morphogenesis between HH stages 8 and 10.

Conducting time lapse experiments of electroporated embryos revealed a highly complex, dynamic coordination of cell movements. Some aspects of these experiments confirm prior findings in the literature through direct observation of endoderm cell movements, while others provided new and unexpected insights. For example, polarization of endoderm cells to align with the AIP lip has been reported (Kidokoro et al. 2018) in fixed samples. However, live imaging revealed a remarkably rapid onset of polarization, with cells switching from disorganized, rounded cells to elongated cells, oriented parallel to the AIP lip in the course of approximately 30 minutes (Figure 5A). Consistent with prior studies of tissue deformations (Shi, Varner, and Taber 2015; Hack et al. 2023) that indicated contraction or compaction of endoderm along the AIP lip, we observed that as cells polarize, they begin crowding toward the midline of the AIP (Figure 5B and Supplemental Movie 1). This tissue-scale shortening was accompanied by intercalation of cells parallel to the AIP lip, suggesting that coordinated cell reorganization, rather than simply acto-myosin contractility on its own, may play a driving role in tissue shortening along that AIP.

**Figure 5:**
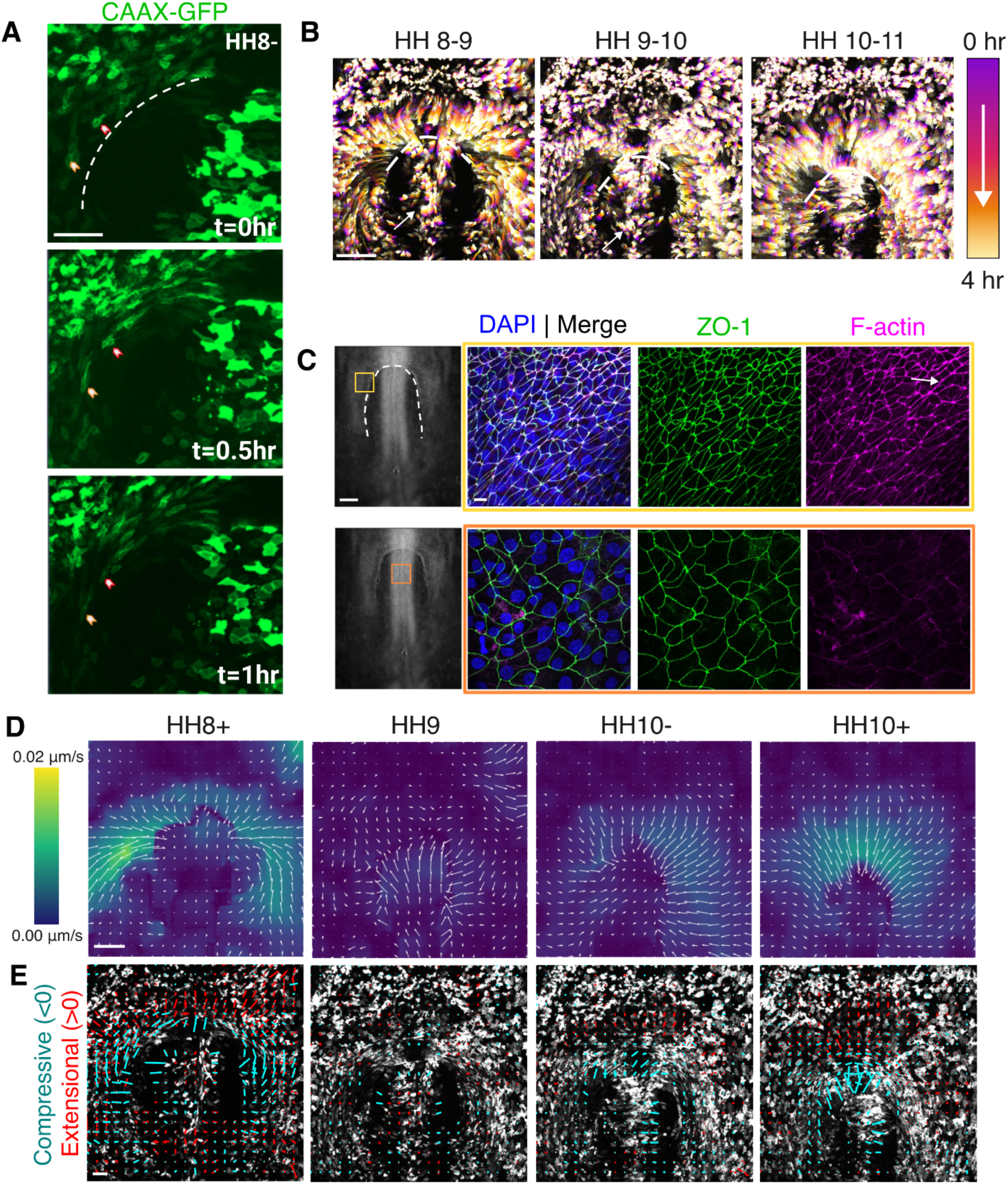
Cell movements, morphology, and tissue deformations at the AIP during foregut elongation. (**A**) Snapshots of a time lapse (Movie S1) dynamic changes in cell morphology of CAAX-EGFP electroporated endoderm cells at the AIP lip (dashed line) at HH stage 8-; red and yellow arrows indicate rapid polarization of cells; scale = 50 μm (**B**) Progressive time-projected cell tracks from time lapse (Movie S2) of a CAAX-GFP electroporated embryo spanning HH stages 8-11; white arrows indicate anterior cell flow in the presumptive foregut roof endoderm. Scale = 200 μm. (**C**) Whole-mount fluorescence imaging of tight junctions (ZO-1, green) and F-actin (Phalloidin, magenta), counterstained for nuclei (DAPI, blue), in the AIP endoderm (top, yellow box) and presumptive roof endoderm (bottom, orange box). White arrow indicates multicellular F-actin cables; scale =10 μm, brightfield image scale = 200 μm. (**D**) Velocity fields quantified from time lapse (Movie S2) of cell movements between HH stages 8+ and 10+. Direction of the velocity vector is indicated by the direction of arrows, and magnitude indicated by both the length of arrows and the underlying heat map (n = 3); scale = 200 μm. (**E**) Principal strain rates computed from velocity fields in (**D**), where major and minor cruciform axes represent maximum and minimum principal strain rates D_1_ and D_2_, respectively, and their orientations represent the associated principal directions. Negative/compressive principal strain rates are indicated in cyan, and positive/extensional strain rates indicated in red; strain rate scale (white) = 0.01%/s, spatial scale is as indicated in **(D)**.

Because much of the focus of prior studies has been on the AIP itself, we were surprised to observe a previously unreported, long-range movement of a posterior population of cells through the AIP along the presumptive roof (Figure 5B, Supplemental Movies S1 and S2), concomitant with AIP descent. This posterior population that appears to contribute to dorsal foregut tube consisted of large, rounded cells with limited apical F-actin enrichment, compared to cells of the AIP lip that were much smaller in area, elongated, and containing cortically localized multicellular F-actin cables (Figure 5C). While the specific role of posterior cells entering the foregut roof is not yet known, these results do further suggest that the model of foregut tube formation as a simple consequence of AIP descensus is incomplete if not incorrect.

To study the biomechanics of cell movements surrounding the forming foregut, we next used particle image velocimetry to quantify tissue flows and deformation rates from time lapse experiments. Velocity fields (Figure 5D) were used to calculate principal strain rates at distinct stages during foregut tube formation. Strain rate refers to the rate of tissue deformation, with positive and negative numbers indicating elongation and shortening rates, respectively. Deformation rates in the AIP endoderm were highly dynamic, with largest magnitudes observed early in tube formation at HH stage 8+ (Figure 5F), correlating with the highest AIP velocity measured (Figure 1B). The orientation of the second principal strain rate (indicating the largest shortening rate when this number is negative) was closely aligned with the contour of the AIP, with largest shortening rates coinciding with regions where cells were most elongated (Figure 5F). While this orientation relative to the AIP persisted, the overall magnitude of shortening diminished significantly from HH stage 9 onwards. These data are consistent with an early role for AIP endoderm cells in shortening the AIP through cell reorganization during the initial phase of foregut tube formation. Consistent with prior studies (Hack et al. 2023), large extensional strains were observed in the extraembryonic tissue immediately anterior to the AIP lip (Figure 5F), likely a result of passive stretch due to pulling forces from the AIP lip, where primarily shortening strain rates were observed. Compared to the AIP, lower strain rates were observed in the presumptive roof, even as cells were displaced anteriorly through the AIP. Together, these analyses further support a central biomechanical role for AIP endoderm in the initial stages of foregut elongation, but a diminishing role as development proceeds.

## 3. Discussion

The present study combines embryological approaches with live imaging and gene misexpression to study the cellular events underlying formation and early elongation of the foregut tube. We find that foregut elongation occurs over a narrower developmental window than previously appreciated (Figure 1D), and that left-right (Figure 2) and dorso-ventral (Figure 3C) symmetry are broken earlier than thought as well. Through tissue-specific gene misexpression studies (Figure 4) and live imaging (Figure 5), we identified a central biomechanical role for endoderm cells lining the AIP in driving foregut morphogenesis, despite finding that these cells do not themselves directly contribute to the elongating tube (Figure 3). Instead, we propose that AIP endoderm produces important extrinsic tensile forces that instruct intrinsic cell responses within the nascent foregut tube to drive its elongation and 3D morphogenesis.

Cell behaviors in the nascent avian foregut are poorly understood, in part due to limitations in microscopy that have precluded high resolution live imaging of cells internal to the foregut. 3D volumetric imaging in the present study does provide some insights into tissue-scale behaviors, however. For example, rapid elongation of the tube between HH stages 8 and 10 involves considerable medio-lateral narrowing (Figure 1E, Figure 3A). This is consistent with convergent extension, an important cell behavior during gut tube formation in zebrafish (Balaraju et al. 2021). While DiI labeling experiments indicated that there is minimal rolling over from the roof to the floor of the tube (Figure 3B), antero-posterior position of labeled cells along the roof did change significantly, consistent with considerable cell rearrangements within this region. By HH stage 11, asymmetries were evident in the foregut tube, including non-uniform cross sections along the antero-posterior axis, and an unexpected expansion on the right side of the tube (Figure 2A). This suggests that the morphological groundwork for later organogenesis begins to appear quite early in the foregut tube. Recent single cell RNA Sequencing work in the E 8.5 mouse embryo corroborates these findings, revealing dorso-ventral and antero-posterior identity already evident at the transcriptional level (Han et al. 2020).

The long range movement of the AIP (Figure 1A) has long been appreciated as a key aspect of foregut elongation, but its specific role has been disputed in the literature. It has been argued, for example, that movement of the AIP and elongation of the foregut roof occur in a wave of lateral-to-medial folding that progresses from anterior to posterior, bringing margins of the definitive endoderm to meet at the midline and zipper the endoderm shut (Bellairs 1953).

Several studies have contested this, suggesting instead that movement of the AIP pulls the floor posteriorly to elongate the tube (Stalsberg and DeHaan 1968; Rosenquist 1971; Varner and Taber 2012; Shi, Varner, and Taber 2015; Hosseini, Garcia, and Taber 2017). The present study conclusively answers this question by way of live imaging, where we observed endoderm cells along the AIP move with the AIP as it descends posteriorly, refuting the zippering hypothesis. We further demonstrate the biomechanical importance of endoderm cells at the AIP through quantification of tissue deformations (Figure 5E and Supplemental Movie 1), and observing that elongation of the foregut is dramatically reduced upon targeted disruption of F-actin in the AIP endoderm (Figure 4C). How these pulling forces translate to elongation of the foregut to triple its original length between HH stages 8 and 10, however, remains an open question; how does the foregut floor sustain such rapid growth intrinsically, without significant addition of cells across the AIP boundary? Although we observe anterior ward flows of cells through the AIP into the forming roof of the foregut tube (Figure 5B), these cells do not roll over the blind end of the tube from roof to floor (Figure 3A). Elongation of the foregut floor must therefore primarily occur either through growth or through insertion of new cells into the floor from lateral regions of the foregut tube.

Fate mapping studies initiated earlier, at HH stage 4, have demonstrated that ventral tissue of the prechordal plate gives rise to descendents that are distributed along the midline of the foregut floor (Kirby et al. 2003). These results suggest that perhaps the prechordal plate is partitioned antero-posteriorly into ventral and dorsal compartments during the initiation of the AIP, and the ventral subpopulation is responsible for elongation of the midline foregut floor through proliferative growth. Broad inhibition of cell proliferation throughout the embryo slows foregut elongation (Hosseini, Garcia, and Taber 2017), supporting the role of cell division in elongation of the foregut. Because fate maps in the present study focused primarily on cells on the AIP lip (Figure 3A), this prechordal plate-derived population was not directly studied here. Curiously, however, we did consistently observe that the ventral midline of the AIP would not express reporters when electroporated with either DNA or RNA (Supplementary Figure S6).

Whether this represents a distinct prechordal plate-derived tissue and why it cannot be labeled using these techniques remains unclear. However, prior fate mapping studies in the mouse have identified this region, the ventral midline of the endoderm lip (VMEL), as a progenitor population giving rise to ventral thyroid, liver, and pancreas (Tremblay and Zaret 2005; Angelo, Guerrero-Zayas, and Tremblay 2012). Consistent with this, we have observed that this cell population does eventually internalize at HH stage 11 (Supplemental Movie 1). However, this is after elongation of the foregut has plateaued (Figure 1D), suggesting that the VMEL does not contribute to initial formation and elongation of the nascent foregut tube.

Because the AIP endoderm has been attributed with both mechanical (Varner and Taber 2012; Hosseini, Garcia, and Taber 2017) and patterning roles (Schultheiss, Xydas, and Lassar 1995; Alsan and Schultheiss 2002) in heart development, it is not yet clear whether the cardiac defects observed upon misexpression of DeAct in the AIP endoderm is due to a loss of contractility alone: severe disruption of the endoderm with DeAct RNA electroporation in many cases resulted in delamination of AIP endoderm cells, potentially diminishing or disrupting the putative organizing role of this tissue for heart development (Anderson et al. 2016). The loss of epithelial continuity with DeAct electroporation, without total removal of the endoderm, suggests that this perturbation may have a more pronounced effect on the mechanical role of AIP endoderm than on its signaling role. Our results are generally consistent with knockout studies in mice where deletion of key endoderm genes such as *Sox17* and *GATA4* produced cardia bifida, and in chick, where laser ablation of lateral AIP endoderm (Kirby et al. 2003) disrupted dextral rotation during cardiac c-looping. Notably, ablation of cardiac progenitors from the epiblast produces a largely normal foregut, despite failure to form a heart, suggesting that reciprocal interactions from the precardiac mesoderm to guide foregut formation are likely limited (Ehrman and Yutzey 1999). Additional studies are needed to understand the mechanism underlying the cardiac phenotypes observed in the present study upon tissue-specific disruption of the endoderm.

In contrast to neurulation, during which an initially flat epithelial sheet undergoes lateral to medial folding to enclose a lumen in the resulting neural tube, gut tube formation among amniote embryos appears to follow distinct physical mechanisms when forming each of three antero-posterior segments, despite resulting in a single continuous epithelial tube. While foregut morphogenesis relies on movement of a portal to elongate the tube, the hindgut forms due to collective movements of posterior endoderm through a stationary portal (Nerurkar et al. 2019), while the midgut tube forms through lateral-to-medial folding events similar to neurulation (Miller et al. 1999). How these processes are coordinated over multiple length and time scales to form the gut as a continuous epithelial tube running the antero-posterior length of the embryo (millimeters long in the HH stage 18 chick embryo) is not known. Further, despite conserved function of gut-derived organs, amniotes and anamniotes form the gut tube by distinct mechanisms as well. Fish and frogs rely on radial intercalation to cavitate a condensed rod of endoderm into a hollow tube (Reed et al. 2009; Chalmers and Slack 2000). As we understand more about foregut and hindgut tube morphogenesis, it may be possible to begin understanding why amniotes evolved such a complex and unique method of tube morphogenesis for the gut, as well as how the disparate events of foregut, midgut, and hindgut development are coordinated. For example, collective movement of roof cells through the AIP in the present study mirrors the collective cell movements through the CIP which form the hindgut. In the hindgut, these movements are organized by a long range FGF gradient (Nerurkar et al. 2019), while broad pharmacologic disruption of FGF signaling disrupts foregut elongation (Hack et al. 2023), suggesting a potential molecular and cellular connection between the mechanisms of foregut and hindgut tube formation. Whether the molecular mechanisms underlying foregut and hindgut morphogenesis are similar is not yet known, but elucidating the signaling and cellular mechanisms that underlie coordination between anterior movement of roof cells, descensus of the AIP, and growth of the foregut floor may provide new insights into the evolution of gut tube formation and broader questions of long range coordination of epithelial morphogenesis in vertebrate development.

## 4. METHODS

### 4.1 Chick embryo collection and ex ovo culture

Fertilized White Leghorn chicken eggs (University of Connecticut Poultry Farm) were incubated at 37°C and approximately 60% humidity until the desired stage. Embryos were harvested on filter paper rings, cultured according to the EC method (Chapman et al. 2001), and staged according to the Hamburger and Hamilton staging system (Hamburger and Hamilton 1951). Embryos were incubated with the dorsal side of the embryo contacting the semisolid culture medium until the desired stage.

### 4.2 RNA and Plasmid DNA Electroporation

Endoderm-specific *ex ovo* electroporation of plasmid DNA was performed as previously described (Nerurkar et al. 2019). Briefly, embryos were placed in a PBS bath atop a 2mm square electrode and electroporation solution consisting of 5 µg/µL plasmid DNA, 5% sucrose and 0.1% Fast Green Dye in PBS, was delivered directly onto the anterior endoderm of HH stage 7-8 embryos. Electroporation was performed by placement of a second 2 mm square electrode approximately 4 mm away from the ventral surface of the embryo, followed by a sequence of five 35V poring pulses of 0.2 msec duration separated by 50 msec with a 10% decay between each pulse, preceding a sequence of five 4V transfer pulses of 5 msec duration separated by 50 msec and 40% decay between each pulse (Nepa 21 transfection system, Nepa Gene, Ichikawa City, Japan). Following electroporation, embryos were returned to EC culture and incubated for 4-5 hours prior to imaging.

While mosaic expression of fluorescent proteins was ideal for visualizing tissue deformations and cell behaviors during foregut formation, functional studies relying on gene misexpression to perturb cell behavior required higher efficiency electroporation along the AIP lip. To achieve this, we turned to RNA electroporation, which produces more uniform and faster expression of proteins than conventional plasmid electroporation (Tran, Dave, and Lansford 2019). Fusion protein DeAct-SpvB-EGFP (Addgene #89446) was subcloned to pCS2 (isolated from pCS2-3nls-EGFP plasmid, Addgene #165400). Linearized DeAct-SpvB-EGFP and pCS2-3nls-EGFP plasmids were used for in vitro transcription of mRNA utilizing SP6 RNA Polymerase (SP6 mMessage Mmachine in vitro transcription kit, Thermo Fisher, #AM1340) , followed by purification via LiCl precipitation (Tran, Dave, and Lansford 2019). 500 ng/µL mRNA with 10% sucrose, and 0.1% fast green in RNAse-free ddH20 was electroporated as above, but with 0% decay of the poring pulse, and the transfer pulse replaced by a second sequence of poring pulses performed with the upper electrode approximately 8 mm from the embryo. GFP signal in endoderm cells was visualized following incubation at 37°C as early as 1 hour post-electroporation. For experiments quantifying foregut elongation and heart tube formation, embryos were imaged using a ZEISS AxioZoom v16 immediately following electroporation, screened for electroporation efficiency and viability following 2 hours of incubation, and placed back into the incubator for 8 hours. Embryos were imaged again following 10 total hours of incubation, and fixed for whole mount immunostaining and imaging. All images were taken at 40x magnification.

### 4.3 Embryology, vital dye labeling, and morphometric analyses

Foregut length was measured by injection of dye into the foregut lumen, followed by bright field imaging. Specifically, foregut lumens were injected with 1% Fast Green dye through the AIP using a pulled glass needle bent to a 45° angle, and immediately imaged to minimize diffusion of the dye from the foregut lumen. Foregut lengths were measured as the distance from the anterior blind end of the foregut tube to the medial point of the AIP, and foregut width was measured at the widest extent of the foregut tube at each stage. Foregut length measurements were normalized by the head-to-tail length of each embryo (the distance from the anterior neuropore to the center of the tailbud), and foregut aspect ratio quantified as the maximum foregut width divided by foregut length. The role of axis elongation in foregut elongation was tested by using a tungsten needle to introduce a cut through all three germ layers at the 4th somite level of HH stage 8 embryos. Dissections were performed sufficiently wide (∼300 µm wide) to prevent tissue healing during the following 10 hour incubation at 37°C to HH stage 10.

Injection of lipophilic vital dyes DiI and DiO (2.5 mg/mL in Dimethylformamide, Invitrogen) was used to label and track tissue movements over time (Nerurkar et al. 2019). A pneumatic microinjector (World Precision Instruments, PV820) was used to inject small volumes of dye through pulled capillary needles. For measurement of AIP displacement, DiI injection of somites was used to mark a local reference point; the relatively static position of somites during foregut elongation was confirmed via live imaging of an HH stage 8 embryo (Figure S1).

### 4.4 Fluorescence microscopy, immunostaining, and EdU staining

For 3-D imaging of the foregut tube, embryos were fixed with 4% paraformaldehyde (PFA) overnight, permeabilized with 0.1% Triton in phosphate-buffered saline (PBS) before incubation in 1ug/mL DAPI (Invitrogen, D1306) at 4°C overnight. Stained embryos were washed for 10 minutes with 0.1% Triton in PBS, followed by two 5 minute washes in PBS. RapiClear 1.49 (SunJin Lab) solution was warmed at 37°C for 30 minutes, then stained embryos were transferred into warmed RapiClear and incubated overnight at room temperature. Cleared embryos were mounted in RapiClear for confocal imaging on a ZEISS LSM 880 confocal microscope with a 40X water immersion objective. Z-stacks spanning the dorso-ventral extent of the foregut were collected with a slice thickness of 0.5 μm. 3D volumes were reconstructed and the foregut lumen was manually traced on sequential transverse optical sections to quantify volumes and surface areas using Imaris (Bitplane).

Section immunofluorescence was carried out as described previously (Nerurkar et al. 2019). Briefly, embryos were fixed overnight at 4°C in 4% PFA, and following a sucrose gradient to 30% in PBS, were embedded in OCT and cryosectioned (Lecia) to 16µm thick slices. Sections were permeabilized with 0.1% Triton in PBS and blocked for 2 hours at room temperature with 5% heat inactivated goat serum, 1% BSA, and 1% DMSO in 0.1% Triton in PBS. Embryos were incubated in anti-E-cadherin (1:200, BD Bioscience 610181) and anti-Laminin (1:200, Abcam ab2034) antibodies in blocking buffer at 4° C overnight, and detected using fluorescently-conjugated secondary antibodies (1:500). F-actin and nuclei were visualized with AlexaFluor 555-phalloidin (1:300, Cytoskeleton PHDH1A) and DAPI (1:1000, Invitrogen), respectively. Whole-mount immunostaining was performed similarly to sections with some modifications. Embryos were permeabilized in 0.1% Triton X, blocked with 10% heat inactivated goat serum in 0.1% Triton X in PBS for 1 hour, then incubated in anti-ZO-1 (1:250, Invitrogen 33-9100), anti-Laminin (1:200, DSHB 3H11), and chick-anti-GFP (1:1000, Abcam ab13970) overnight at 4° C. Imaging of whole-mounts and sections was conducted on a Zeiss LSM 880 laser scanning confocal microscope with either 40x water or 63x oil immersion objectives.

### 4.5 Time-lapse imaging

HH stage 8- electroporated embryos were transferred to glass bottom dishes (Corning) coated with EC culture medium (Chapman et al. 2001). To improve viability of HH stage 8 embryos cultured dorsal side up on semisolid medium, the original EC culture protocol was adapted by lowering the agar concentration to 1.5mg/mL. Once placed in the glass bottom dish, two thin strips of lens paper were placed on the area opaca lateral to the embryo, in order to minimize drift during image acquisition. Kimwipes dipped in sterile, distilled water were placed in the margins of the glass bottom dish and the lid sealed with parafilm to limit evaporation. Time-lapse images were acquired using an inverted ZEISS LSM880 confocal microscope with a heated chamber. GFP signal was collected using a GaAsp detector at 1024 x 1024 pixel resolution and 5 μm z-intervals, and 5 minute time intervals.

### 4.7 Strain Rate Analysis

A custom image analysis pipeline was developed in Python using the scikit-image library (van der Walt et al. 2014) to calculate strain rates from z-projected time lapse movies. For each time point, two images collected 10 minutes apart were smoothed with a gaussian filter and used to compute 2D velocity fields by optical flow analysis. The iterative Lukas-Kanade optical flow algorithm (Wedel et al. 2009) was applied with a window size of 50 pixels, which is estimated to be about 2 times larger than the average cell size in each image. The velocity field was downsampled using block reduction to reduce the effects of noise resulting from the optical flow registration. The deformation rate tensor (**D**) was computed

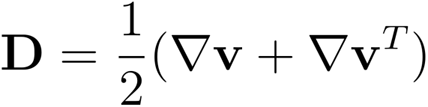

where 𝛻 denotes the gradient operator and **v** is the velocity field computed by optical flow.

Principal deformation rates and orientations were computed by eigendecomposition of **D**. To remove artifacts associated with the disparate focal planes of the AIP lip and presumptive roof in z-projections, an open spline was manually annotated along the AIP of each image prior to strain rate calculation using an interactive matplotlib library. Principal strain rates calculated along the interface between the AIP and the presumptive foregut roof were removed from analysis..

### 4.8 Statistical Methods

Quantitative data are presented as mean plus or minus the standard deviation.

Comparison of means between two groups made with a Student’s t-test. Comparison of means across three or more groups were made using One-way ANOVA, followed by post-hoc Tukey’s test for comparison between groups. Pearson coefficients, and p-values were calculated to determine correlation between two groups (Figure 1F).

### 4.9 Data and Code Availability

Data and data analysis code will be made available upon request.

## Funding sources and Acknowledgements

We thank the members of the Nerurkar lab and the Kasza lab for their scientific input and valuable feedback. In particular, we thank Panagiotis Oikonomou for his assistance with data analysis. This work was funded by the National Institute of General Medical Sciences (R35GM142995 to N.L.N.), with further support from the Columbia University Digestive and Liver Disease Research Center (1P30DK132710).

## CRediT authorship contribution statement

**O. Powell:** Conceptualization, Methodology, Software, Validation, Formal analysis, Investigation, Data Curation, Writing - Original Draft, Review & Editing, Visualization. **E. Garcia:** Validation, Investigation, Writing - Review & Editing. **V. Sriram**: Investigation, Formal analysis, Validation. **Y. Qu:** Investigation, Validation. **N.L. Nerurkar:** Conceptualization, Methodology, Resources, Writing - Original Draft, Writing - Review & Editing, Supervision, Project administration, Funding acquisition.

## SUPPLEMENTAL FIGURES

**Figure S1:**
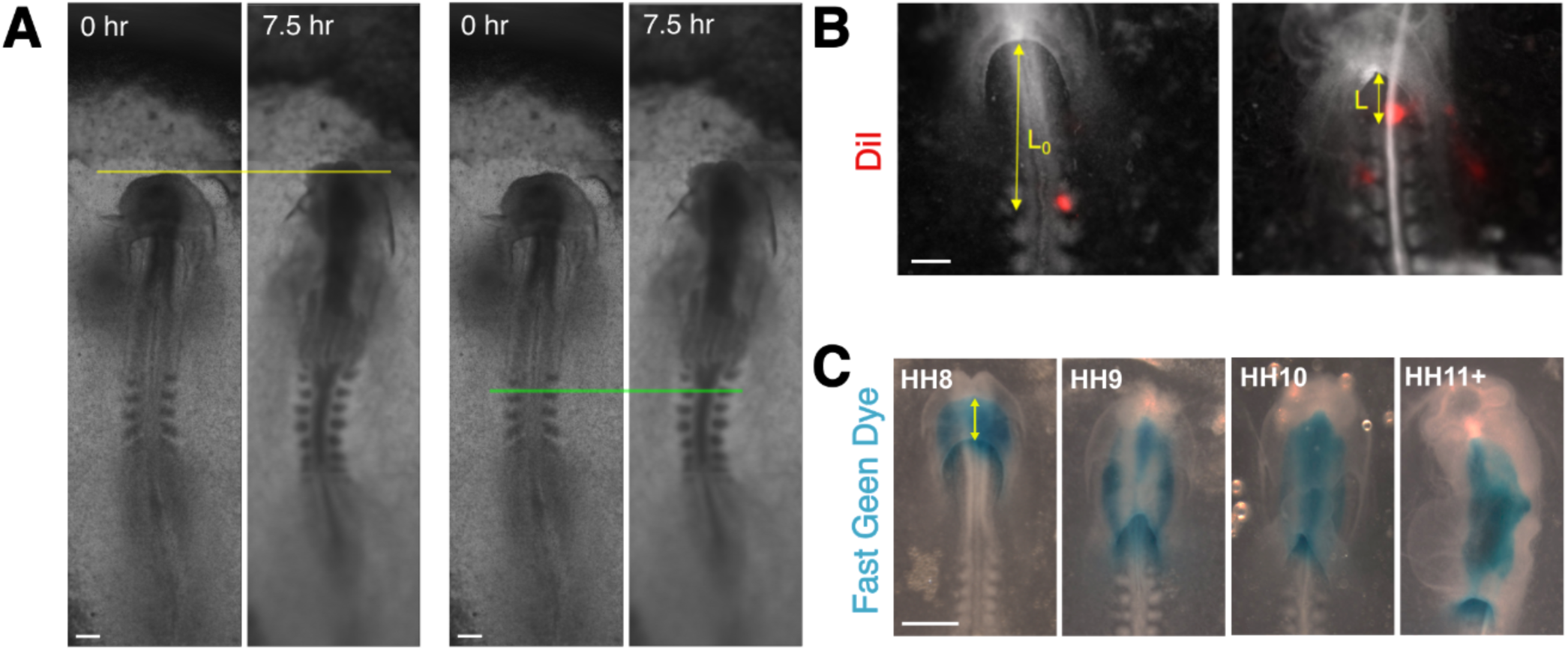
Quantification of AIP displacement and foregut length. (**A**) Yellow and green lines overlaid on snapshots of a brightfield timelapse indicating the initial position of the anterior neuropore (left two panels) and second somite (right two panels), respectively. The anterior neuropore (left), used previously as a reference point for measuring AIP displacement, extends anteriorly by approximately 100 μm during foregut formation. On the other hand, early somites, and somite 2 in particular, remain relatively stationary (Supplemental Movie 2), and were hence used as a reference point for AIP velocity measurements. (**B**) Injection of somites with DiI (red) was used to establish a fixed reference point, against which displacement of the AIP was quantified by change in the distance between the AIP and DiI label (L-L_0_) over intervals spanning HH stages 8-9, 9-10, and 10-11; scale = 200 μm. (**C**) Brightfield time course of lumen visualization by Fast Green injection to quantify foregut length between HH stages 8 and 11+; scale = 500 μm.

**Figure S2.**
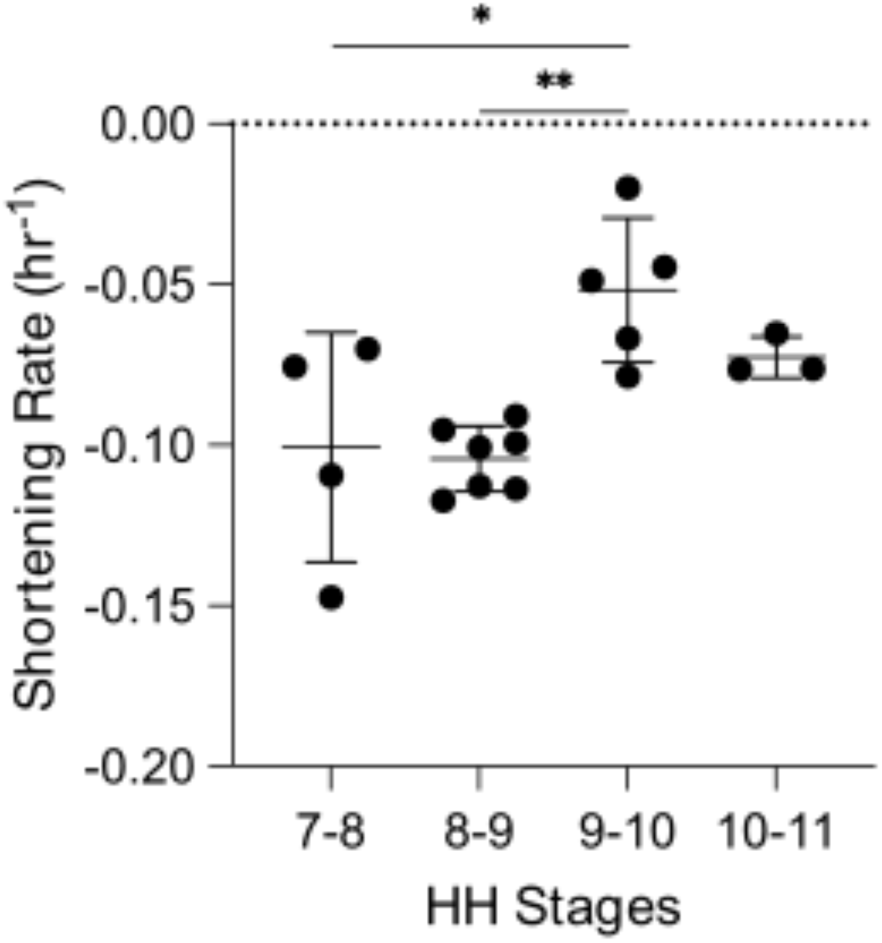
Tissue shortening rate along the AIP coincides with elongation of foregut. Shortening rate of AIP endoderm quantified from movement of DiI labels injected between HH stages 7 and 10; *p<0.05, **p<0.01 by one-way ANOVA followed by Tukey’s post-hoc tests. The ratio between neighboring labels (current length (*l*) normalized by original length (*L*)) was measured using ImageJ and defined as the stretch ratio (*λ*). AIP shortening rate was calculated as described by Shi et al. (2015): 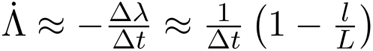

**Figure S3.**
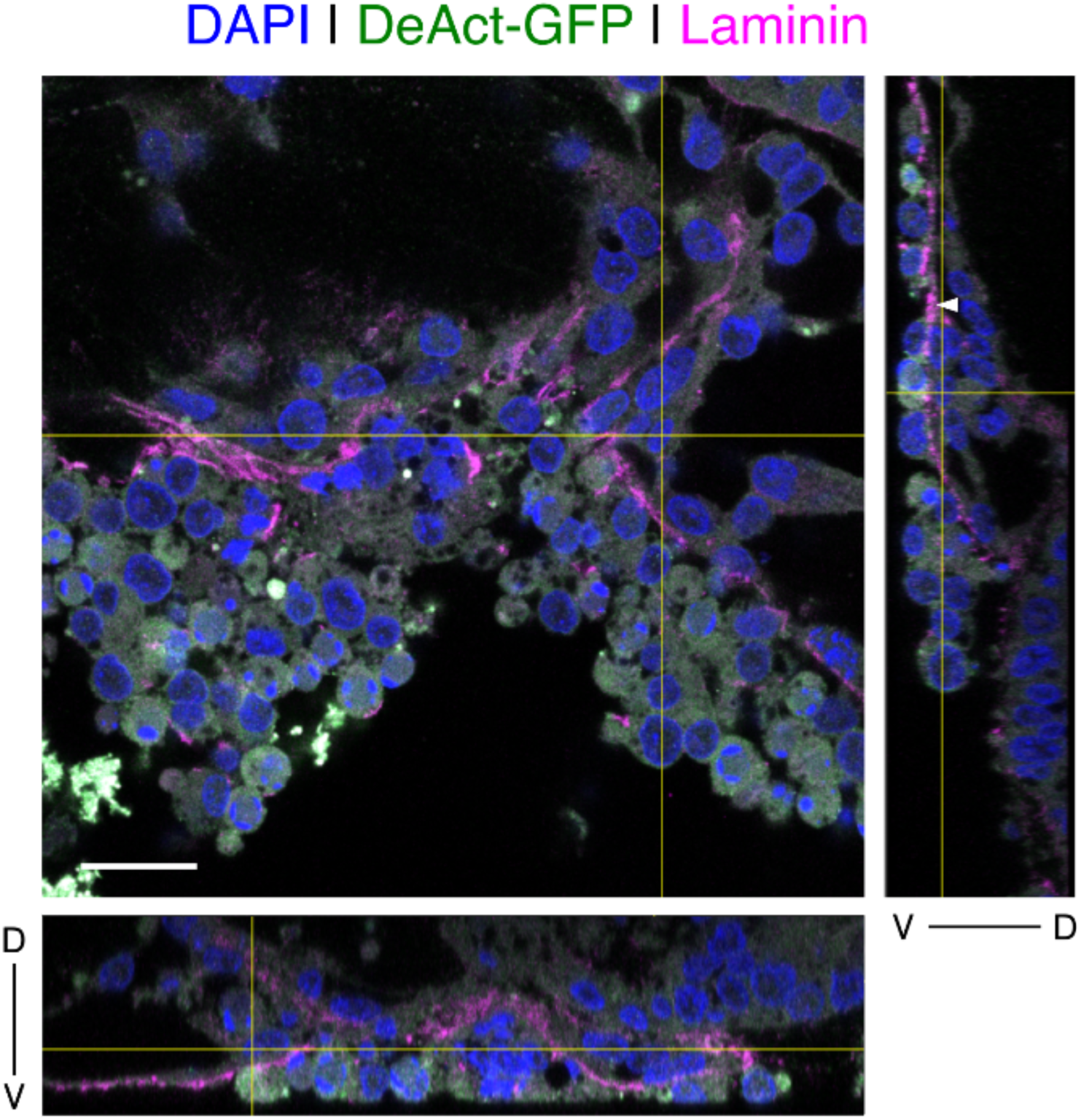
Rounding and delamination of endoderm cells following misexpression of DeAct-GFP. Confocal imaging of AIP region 10 hours after mRNA electroporation of DeAct-GFP, with rounded, GFP-expressing cells, immunostained for laminin (magenta) and counterstained with DAPI (blue). DeAct-GFP electroporated embryos contained highly rounded GFP+ cells (green), and in many cases, gaps in the epithelium likely resulting from delamination of DeAct-GFP expressing cells, leaving behind acellular regions of basement membrane (indicated by white arrowhead in ortho-views); D = dorsal, V = ventral; scale = 50 μm.

**Figure S4.**
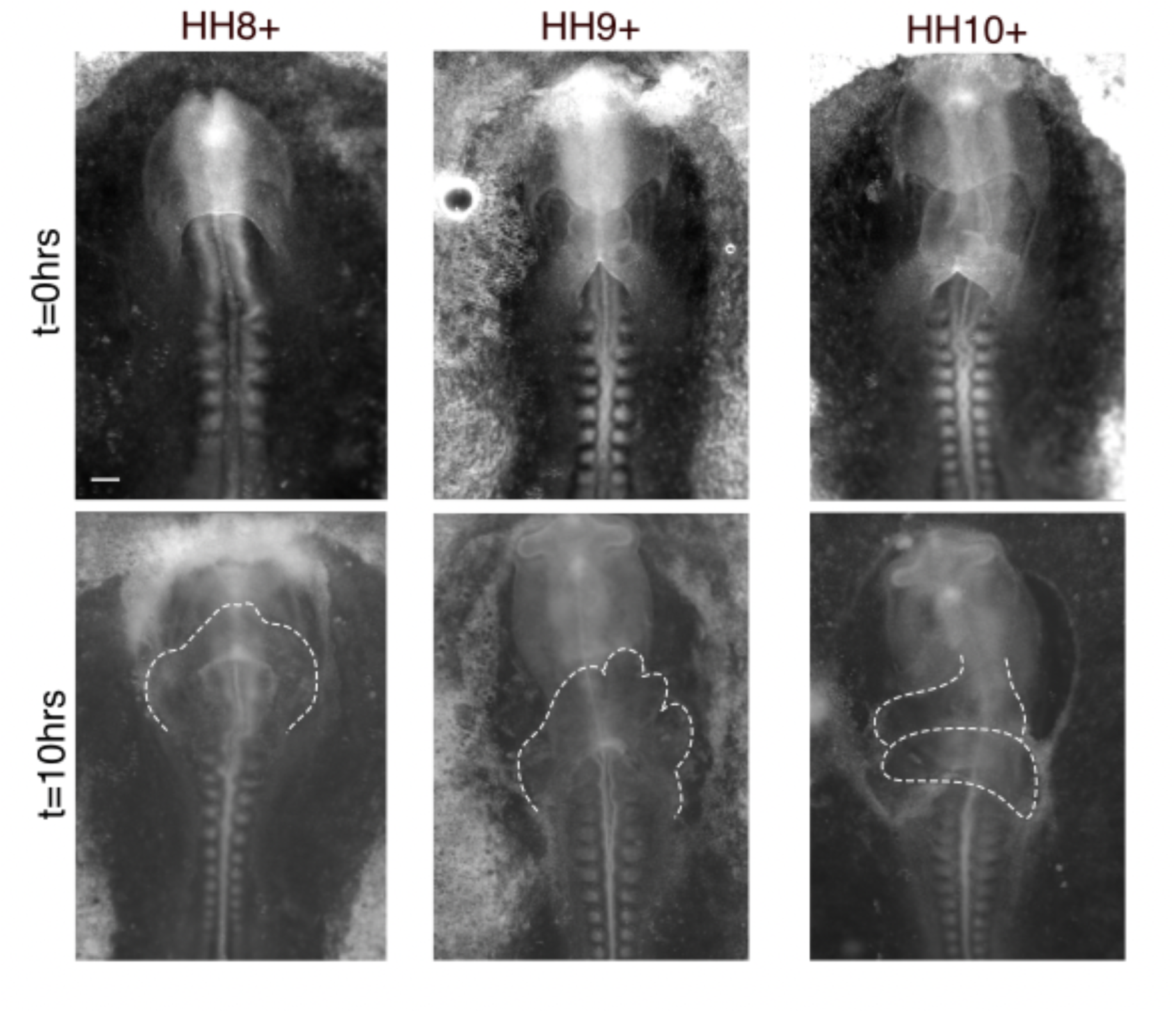
Range of heart defects observed following misexpression of DeAct-GFP in the endoderm is stage dependent. Representative brightfield images of ‘mislooped’ heart tube phenotypes that result from DeAct-GFP electroporation at HH stages 8+ (left), 9+ (middle), and 10+ (right). Embryos electroporated between HH stages 8- to 8+ developed unlooped heart tubes, bilateral heart tubes, or lacked heart tubes entirely following 10 hours of incubation. Embryos electroporated between HH stages 9- to 9+ developed heart tubes with many irregular folds and no particular looping directionality, or developed enlarged heart tubes that remained at the midline of the embryo. Embryos electroporated between HH stages 10- to 10+ developed enlarged heart tubes with a flat morphology and widened omphalomesenteric veins. Scale = 200 μm.

**Figure S6.**
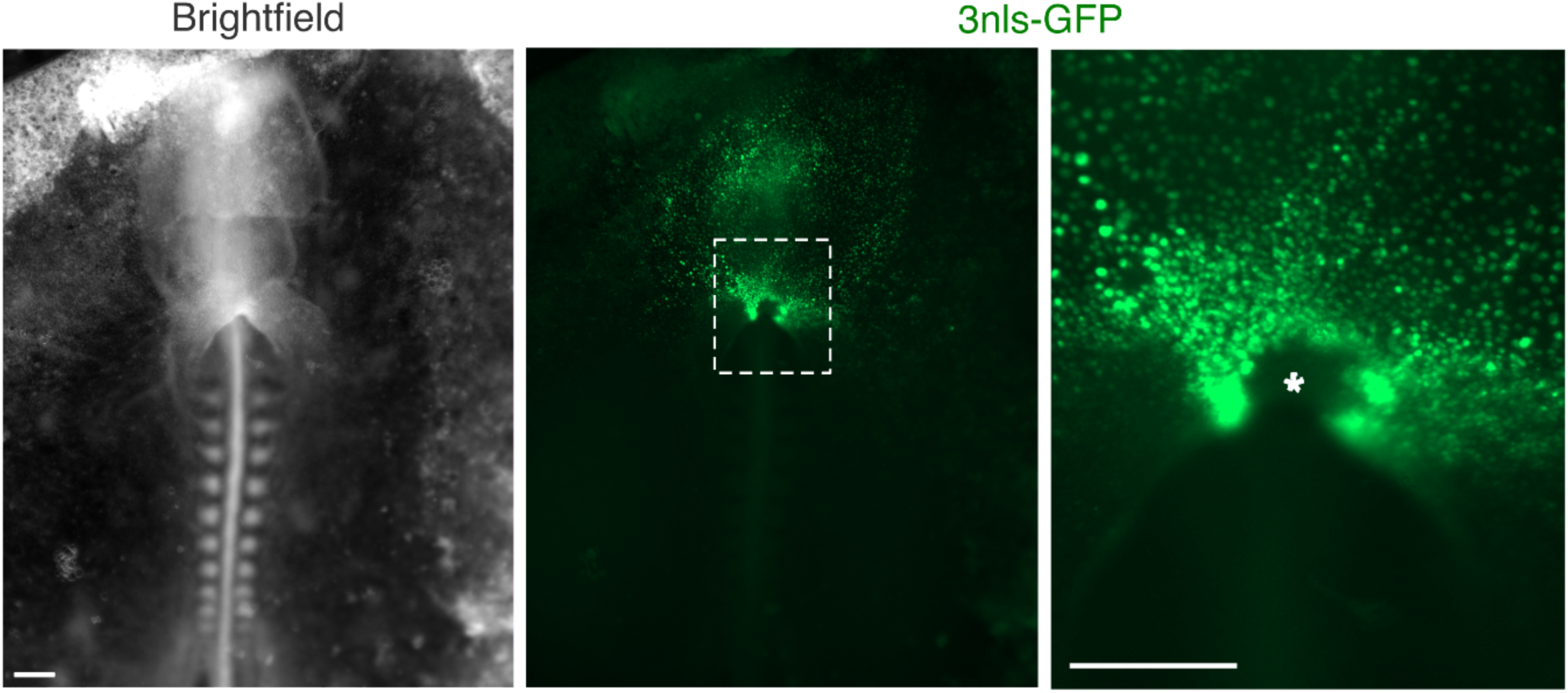
Medial AIP endoderm resists transfection. Representative image of an embryo electroporated with 3nls-GFP at HH stage 8 and incubated for 10 hours. Scale = 200 μm. High-efficiency (>70% of cells) anterior endoderm-targeted electroporation of embryos at HH stage 8 and earlier consistently (n = 8) reveal a cluster of cells located at medial AIP that are not amenable to transfection. *In vivo* live imaging has shown that this medial population of cells eventually involutes into the foregut lumen after foregut elongation has slowed, at HH stage 11 (Movie 1).

